# A Distinct Interphase Microtubule State Marks Host Cell Permissiveness to *Chlamydia pneumoniae* Entry

**DOI:** 10.64898/2026.06.29.735230

**Authors:** Katharina Schenk, Johannes Hegemann, Ursula Fleig

## Abstract

Entry of intracellular pathogenic bacteria is widely considered an actin-driven process, potentially overlooking contributions of the microtubule cytoskeleton. Here, we identify a host microtubule state as a determinant of early infection efficiency by *Chlamydia pneumoniae*. Human cells enriched in acetylated microtubules are preferentially infected, whereas detyrosinated microtubules show no such association. Pharmacological stabilization of microtubules via Taxol enhances infection, while selective elevation of acetylation with Tubacin does not, indicating that microtubule stability rather than acetylation alone is critical. Thus, a pre-existing interphase microtubule architecture supports *C. pneumoniae* entry. Consistently, mitotic cells, characterized by a reorganized microtubule architecture, remain permissive but show severely reduced infection efficiency. In addition, infection induces a dose-dependent increase in microtubule acetylation that requires bacterial viability and is not observed during uptake of *Yersinia pseudotuberculosis* effector protein Invasin-coated beads, indicating that entry/internalization alone is insufficient to trigger this response. To probe how early chlamydial secreted effectors might engage the microtubule cytoskeleton, we focused on the conserved TarP family member CPn0572, an actin and microtubule regulator, which increases microtubule acetylation when ectopically expressed in human cells. Controlled expression of microtubule-localized CPn0572 in the yeast *Schizosaccharomyces pombe* leads to altered microtubule dynamic and mechanical behaviour, promoting force-bearing microtubules. Together, these findings show that distinct interphase microtubules define a permissive cellular state for bacterial entry and suggest that early chlamydial effector activities might promote a specific microtubule persistence phenotype.

## Introduction

The host cytoskeleton is a central target of bacterial pathogens, which exploit actin filaments and microtubules (MTs) to promote entry, intracellular trafficking, and replication and exit [1–3]. Remodelling of the actin cytoskeleton is widely recognized as the primary driver of bacterial internalization, as many invasive bacteria induce localized actin polymerization to promote membrane deformation and uptake. This paradigm is well established for intracellular bacterial pathogens, including the *Chlamydiae*, where early type III-secreted effectors such as TarP family members remodel cortical actin to facilitate entry and co-localize as early as 15 minutes post-infection with actin [4–7].

In contrast, the contribution of MTs to bacterial entry remains poorly defined and is supported primarily by indirect evidence. Early studies reported that pharmacological disruption of MTs reduces bacterial internalization, suggesting a role in invasion; however, these studies rely on global MT cytoskeletal perturbation and do not resolve if MT contribute to the entry process [8–10].

Consistently, MTs have been predominantly assigned functions during intracellular trafficking, perinuclear positioning, and organization of pathogen-containing compartments, as shown for dynein-dependent transport of nascent *Chlamydia* inclusions and effector-driven reorganization of MTs at the inclusion surface [11, 12]. Even in systems where MT-dependent processes act immediately after uptake, such as dynein-dependent membrane uncoating during *Shigella flexneri* vacuolar escape, these findings do not establish a direct role for MTs in bacterial entry [13].

Thus, whether and how MTs influence bacterial internalization remains unresolved. This is particularly relevant given that MTs have key roles in a large number of cellular functions including mechanotransduction and intracellular transport. Their structure enables them to withstand compressive forces and resist tension, while their highly dynamic behaviour allows them to rapidly reorganize in response to intrinsic and extrinsic cues. Due to their association with MT-associated proteins [14] and the existence of the so-called “tubulin code”, the function and properties of specific MTs α-β-tubulin filaments within a single cell can be highly variable. The tubulin code consists of tubulin isotypes and a variety of posttranslational modifications (PTMs) [15]. A subset of PTMs is mainly found on specific MTs such as stable MTs. For example, the PTM Lys40 of α-tubulin acetylation is a consequence of stabilized MTs and results in long-lived MTs that can withstand mechanical stresses due to increased MT filament flexibility, while the PTM detyrosination i.e. removal of the C-terminal tyrosine from α-tubulin also mainly occurs on stable MTs with properties that alter binding of MT motors and microtubule-associated proteins [16, 17].

These two types of modification have been found in the cage-like MT network surrounding established chlamydial inclusions [18, 19], but it is unclear if such MT modifications might have an impact on the ability of a bacterial pathogen to enter a host cell.

This question is the focus of our analysis using infection by C*hlamydia pneumoniae* elementary bodies (EBs) as a test system. *Chlamydiae* are obligate intracellular bacterial pathogens responsible for a significant number of serious human diseases. They share a conserved biphasic developmental cycle that alternates between infectious EBs and intracellular, replicative reticulate bodies (RBs), with EB entry representing a critical step that determines successful infection. Members of the genus include important human pathogens such as *Chlamydia trachomatis*, a leading cause of bacterial sexually transmitted infections and preventable blindness, and *Chlamydia pneumoniae*, a respiratory pathogen associated with community-acquired pneumonia and chronic inflammatory conditions [20, 21].

Consistent with a functional interaction between *Chlamydia* and the host MT cytoskeleton, multiple chlamydial proteins have been shown to modulate MT organization, and depolymerization of MTs reduces infection efficiency [12, 22–27]. In *C. pneumoniae*, the conserved TarP-family early effector CPn0572 directly associates with and remodels both actin and MTs [7, 28, 29], highlighting the capacity of secreted effectors to engage both cytoskeletal systems. Together, these observations suggest that targeting of the MT cytoskeleton is a coordinated aspect of chlamydial infection.

Here, we address the importance of host MTs in bacterial entry/early infection. We show that specific PTM acetylated interphase MTs define a highly permissive cellular state for bacterial entry while another interphase MT PTM namely detyrosination or the mitotic cell cycle stage with an altered MT architecture do not. Additionally, EB uptake itself leads to an increase in MT acetylation in a viability-dependent manner suggesting entry-linked MT modulation via chlamydial early effector proteins. Analysis of the controlled expression of the early effector protein *C. pneumoniae* TarP in a yeast test system shows that CPn0572 has an effect on MT stability as it uncouples MT plus-ends from normal membrane-induced catastrophe responses.

## Results

### Increased infection efficiency of *C. pneumoniae* EBs correlates with number of acetylated host cell MTs

We had shown previously that drug-induced depolymerization of the interphase MT cytoskeleton prior an infection with *C. pneumoniae* EBs followed by re-polymerization of the MTs during infection resulted in a significant reduction of *C. pneumoniae*-infected epithelial cells [27]. However, as we scored infection after 30 hrs of incubation it was unclear if MTs played a part in entry/early infection process. Synchronization of *Chlamydia* EB entry is generally accomplished by allowing bacteria to bind host cells at 4°C and then shift to 37°C for rapid internalization [30]. Incubation of cells at 4°C leads to a fast loss of most MT polymers [31] and thus EB host cell entry is generally carried out under MT defective conditions. To synchronize EB entry but also have the possibility to determine the impact of the MT cytoskeleton on entry/early infection, our rationale was to use the established infection protocol but determine the ability of EBs to infect human epithelial U2OS cells with different amounts of MT PTMs. We analysed the involvement of acetylated and detyrosinated MTs, as both are described as markers for stable MTs [32]. Furthermore, both PTMs have been previously shown to surround the chlamydial inclusion in a MT-cage-like manner in later infection [18]. The experimental set-up is shown in Fig 1A: U2OS cells were incubated on ice for 30 min prior to *C. pneumoniae* infection. This treatment resulted in loss of the MT cytoskeleton with only a small number of acetylated and detyrosinated MTs still present (Fig 1A, middle panels, Fig S1A-C). After EB addition at 4°C, cells were incubated at 37°C for 10 minutes to allow MT re-polymerization and EB internalization (Fig 1A). We chose 10 min post infection (pi) as the time point for our analysis, as at this time point MTs are partially re-polymerized with some defined longer filaments (Fig 1A, right most panels). As cells in a population have variable amounts of PTM MTs resulting in cells with different amounts of MT subpopulations, we could determine if specific MT subpopulations such as detyrosinated or acetylated MTs altered EB infection efficiency (Fig 1B). We added *C. pneumoniae* EBs (MOI 10) to ice-treated U2OS cells, fixed them at the 10-minute time point (Fig 1A) and determined the number of internalized EBs and the percentage of detyrosinated or acetylated MTs versus the entire MT population.

**Fig 1.**
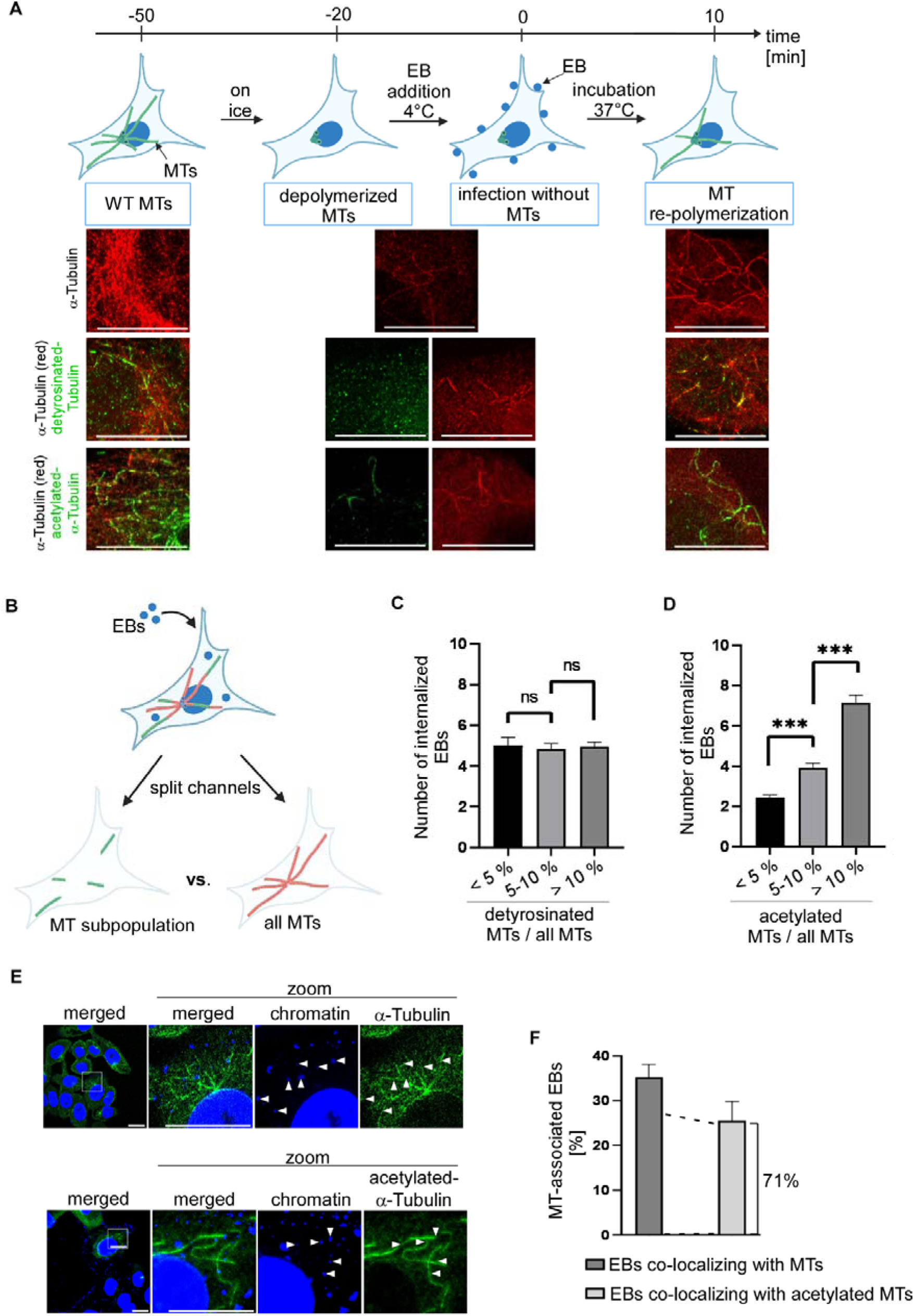
*Chlamydia pneumoniae* infection efficiency depends on MT composition of host cell. **A** Schematic representation of the experimental set-up. Cells were incubated on ice for 30 min to depolymerize MTs. *C. pneumoniae* EBs (MOI 10) were added at 4°C followed by 10 min incubation at 37°C before fixation. Representative confocal fluorescence images of U2OS cells for the various time points. MTs are visualized with an anti-_α_-tubulin antibody (red, all lanes). Detyrosinated MTs are shown in green using an anti-detyrosinated-tubulin antibody (middle lanes). Acetylated MTs are stained with anti-acetylated-_α_-tubulin antibody (green) (bottom lanes). Images are shown as maximum intensity projection. Scale bars: 10 µm. **B** Diagrammic representation of quantification of MT subpopulations in *C. pneumoniae* infected cells and the number of internalized EBs. All MTs are visualized in red with an anti-_α_-tubulin antibody and the MT subpopulation of interest with the respective antibody in green. DNA/EBs are stained with DAPI in blue. ImageJ was used to split the merged image into two single frames: one representing only the MT subpopulation (green) and the other all MTs (red). The ImageJ Coloc2 Plugin was used to determine the amount of the MT subpopulation/ all MTs. **C** Quantification of the number of internalized EBs in cells with different amounts of detyrosinated MTs. Cells were treated as described in **A** and the amount of detyrosinated MTs was determined as in (**B**). n=3 each representing 30 cells. Error bars denote ±SEM, ns; not significant (two-tailed unpaired Student’s *t*-test). <5% = 2,74% ± 1,04%; 5-10% = 8,45% ± 1,7%; >10% = 17% ± 4,86. **D** Quantification of the number of internalized EBs in cells with different amounts of acetylated MTs. Cells were treated and analysed as described in **A** and **B**. n=3 each representing 30 cells. Error bars denote ±SEM, p<0.001 (***) (two-tailed unpaired Student’s *t*-test). <5% = 3,0% ± 1,42%; 5-10% = 7,0% ± 1,12%; >10% = 19,73% ± 9,18%. **E** Representative confocal images of internalized EBs in U2OS cells co-localizing with any type of MTs (top) or acetylated MTs (bottom). Cells were treated as described in **A.** All MTs are visualized via an anti-_α_-tubulin antibody (top) and acetylated MTs with an anti-acetylated-_α_-tubulin antibody (bottom). DNA/EBs are stained with DAPI (blue). Arrow heads show EBs co-localizing with MTs. Images shown are a single layer of the z-stack. Scale bars: 10 µm. **F** Quantification of the number of internalized EBs co-localizing with any MT structure or acetylated MTs. n=3 each representing at least 127 internalized EBs. Error bars denote ±SEM.

Cells with different amounts of detyrosinated MTs were put into 3 categories with detyrosinated MTs/all MTs <5%, 5-10% or >10% (minimum of 2,7% in <5% fraction; maximum of 17% in >10% fraction) and the number of internalized EBs was determined for each category. We found no significant difference in the number of internalized EBs (Fig 1C). Thus, varying amounts of detyrosinated host MTs do not affect EB infection efficiency. Next, we tested if the amount of acetylated MTs in a cell had an effect on EB infection efficiency. We analysed host cells with <5%, 5-10% or >10% acetylated MTs versus total number of MTs (minimum of 3% in <5% fraction; maximum of 19,7% in >10% fraction). Interestingly, there was a direct correlation between the number of acetylated MTs and the number of internalized EBs (Fig 1D). Cells in the <5% fraction had approx. 2,4 EBs, while cells in the >10% fraction had on average 7 EBs (Fig 1D). Thus, cells enriched in acetylated MTs are preferentially infected by *C. pneumoniae* in a dose-dependent manner.

Next, we analysed the subcellular localization of these EBs. At the 10-minute time point (Fig 1A) single MT filaments can be scored easily due to the only partial MT re-polymerization and thus we could determine if EBs were present on MTs. 35% of the internalized EBs co-localized with MTs (Fig 1E top and 1F) and 71% of these EBs co-localized with acetylated MTs (Fig 1E bottom and 1F). As we analysed cells independently of their MT subsets and approximately 9% of MTs were acetylated when considering the entire cell population, these data point to preferential association of EBs with acetylated MTs.

### Long-lived MTs increase the infection efficiency of *C. pneumoniae*

The above analysis showed that the presence of acetylated MTs had a dose-dependent positive impact on the infection efficiency of *C. pneumoniae*. MT acetylation occurs on stable MTs and makes these MTs long-lived [33]. Thus, we asked if MT acetylation *per se* results in more internalized EBs or if the longevity is the cause of the increased infection efficiency. Therefore, U2OS cells were treated with either Taxol or Tubacin. Taxol stabilizes MTs resulting in their acetylation making them long-lived [34], while Tubacin inhibits the tubulin deacetylase resulting in an increased length of the acetylation segments without altering MT stability and longevity [35, 36]. Taxol or Tubacin treated cells showed both a similar and significant increase in the number of acetylated MTs compared to DMSO-treated control cells (Fig 2A, B and Fig S2A). Incubation on ice of Taxol or Tubacin treated cells confirmed that Taxol treatment resulted in stable MTs while Tubacin increased the amount of acetylated MTs without impacting MT stability (Fig S2B). Thus, we infected DMSO (control), Taxol or Tubacin treated cells for 10 min with *C. pneumoniae* EBs and determined the number of internalized EBs via inside-out staining (Fig 2C). Here, non-internalized EBs are stained without prior permeabilization in a first step, followed by permeabilization and a second staining now including internalized EBs [37]. The Tubacin-treated cells showed no significant difference in the number of internalized EBs compared to the control cells (Fig 2D, E). In contrast, we observed significantly more internalized EBs in the Taxol-treated cells compared to the control cells (Fig 2D, E). The ability of EBs to adhere to control cells or Taxol-treated cells were similar (Fig S2C) demonstrating that the internalization process was affected. Thus, MT acetylation *per se* is not the cause of increased infection efficiency, but stable MTs that have been acetylated and thus have altered properties.

**Fig 2.**
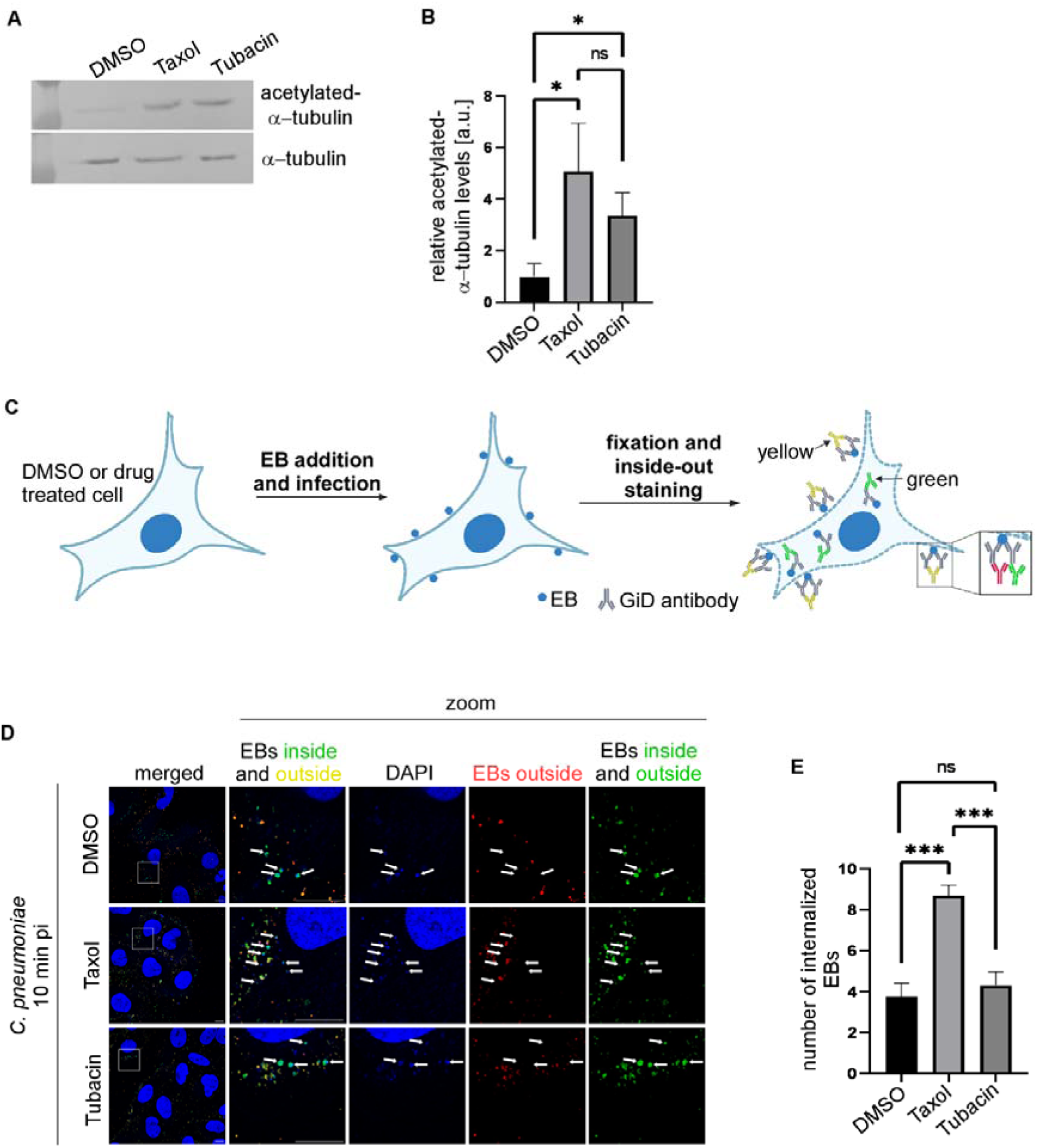
Taxol-treated cells are infected by *C. pneumoniae* with higher efficiency. **A** Western blot analysis of acetylated MTs in differently treated U2OS cells. Cells were treated either with DMSO (0,1%), Taxol (10 µM) or Tubacin (10 µM) for 2 h at 37°C prior to lysis. Samples were separated on a 10% SDS-PAGE followed by Western blot analysis. Western blots were probed with anti-acetylated-_α_-tubulin or _α_-tubulin antibody. The blot shown is a representative of three repeats. **B** Quantification of relative acetylated-_α_-tubulin levels shown in **A**. Band intensities from Western blots of acetylated-_α_-tubulin and _α_-tubulin in cells treated with Taxol or Tubacin were determined. The ratio of acetylated-_α_-tubulin levels to _α_-tubulin levels was analysed and compared relative to control cells (The ratio for DMSO control was set to 1). ImageJ was used for band intensity quantification. n=3 independent experiments, Error bars denote ±SEM, p<0.05 (*), ns; not significant (two-tailed unpaired Student’s *t*-test). **C** Schematic representation of experimental setup: cells were incubated with DMSO, Taxol or Tubacin for 2 h at 37°C followed by *C. pneumoniae* infection (MOI 10) at that temperature. 10 min pi cells were fixed and internalized EBs were visualized with an inside-out staining: Non-internalized EBs were stained with anti-GiD antibody (red) before permeabilization. After permeabilization non-internalized and internalized EBs were visualized via an anti-GiD antibody (green). While non-internalized EBs appear yellow (red and green), internalized EBs are green. **D** Representative confocal images of U2OS cells treated with DMSO, Taxol or Tubacin and infected with *C. pneumoniae* EBs as described in **C**. EBs were stained with an inside-out staining and DNA with DAPI (blue). Images show maximum intensity projection. Arrows mark internalized EBs (green). Scale bars: 10 µm. **E** Quantification of the number of internalized EBs in the differently treated cells shown in **D**. n=3 representing 30 cells each. Error bars denote ±SEM, p<0.001 (***), ns; not significant (two-tailed unpaired Student’s *t*-test).

### Mitotic cells are infected by *C. pneumoniae* with highly attenuated efficiency

Our data show that a specific subset of interphase MTs is important for EB infection efficiency. We next asked, if chlamydial infection was also possible in the absence of the interphase MT array. This scenario occurs when the host cell is in the mitotic phase of the cell cycle. Interphase MTs are dismantled and MTs form the mitotic spindle. Endocytosis is reduced during early mitosis, but reactivated from anaphase onwards [38]. Notably, some receptor-specific internalization pathways remain active despite the general reduction in endocytic activity. This includes EGFR uptake which is, among others, utilized by *C. pneumoniae* for host cell entry [38, 39]. In our non-synchronized U2OS population approximately 5% of cells were in different stages of mitosis. To study more mitotic cells, we synchronized cells prior to infection by treating cells with the CDK1-inhibitor RO-3306. RO-3306 blocks cells at the G_2_/M transition and thus allows synchronized entry into mitosis after the release from the block [40]. In our experimental set-up, 1.5 h after RO-3306 washout approximately 36% of the cells were in mitosis and this number did not change significantly 2 h and 2.5 h after washout (Fig S3A, S3B). The remaining cells were in various non-mitotic stages and infection of those cells was also determined. This subpopulation is termed “non-mitotic” in the text and in Fig 3.

**Fig 3.**
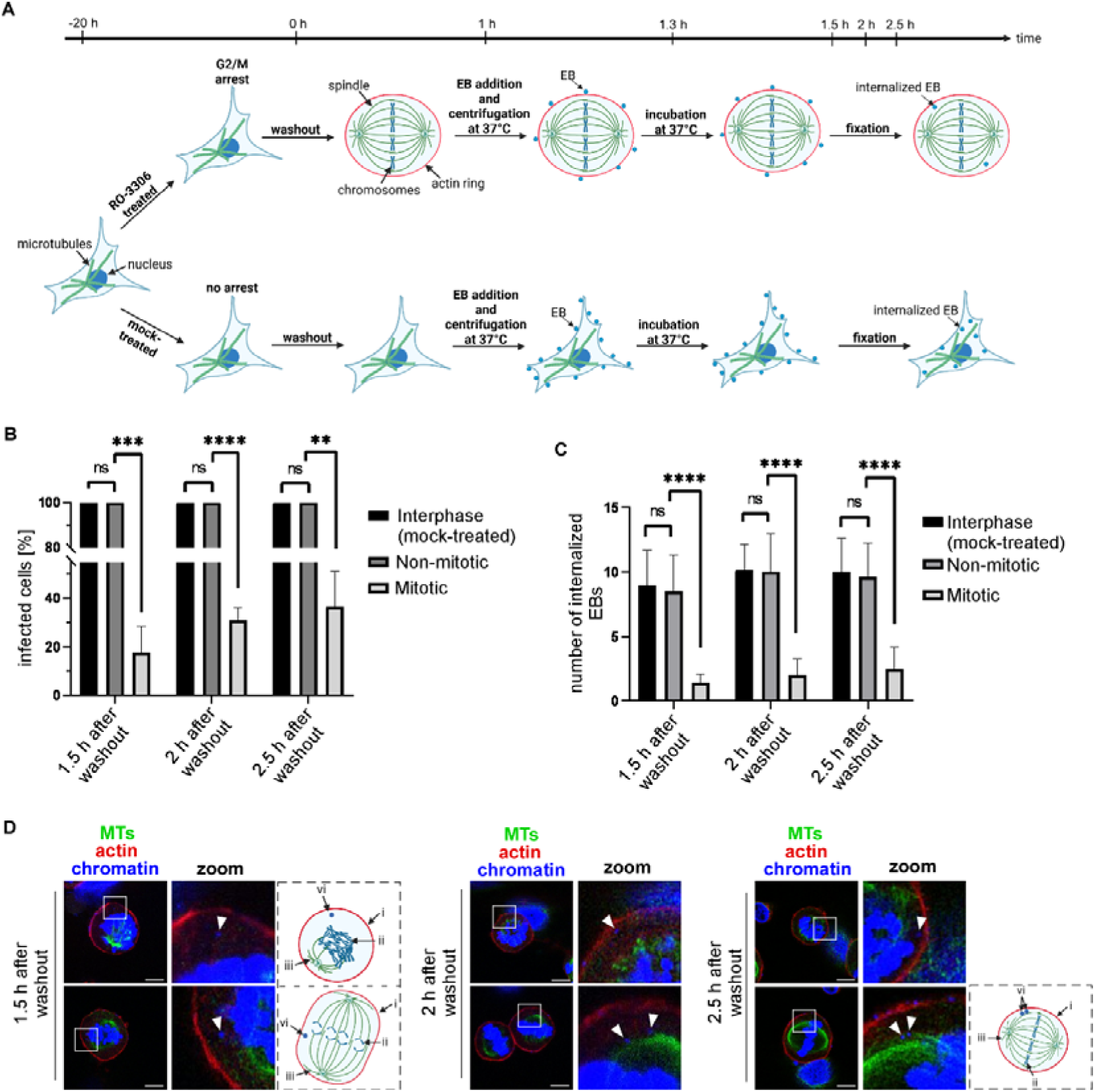
Mitotic cells have a highly reduced capability to be infected by *C. pneumoniae.* **A** Schematic representation of the experimental set-up. U2OS cells were either mock (DMSO)- or drug-treated with 6 µM of the CDK1-inhibitor RO-3306 for 20 hrs. Mock-treated cells were treated as drug-treated cells. DMSO- and RO-3306-treated cells were washed and RO-3306 released cells entered M-phase (mitotic cell shown in cartoon represents all mitotic stages). 1 h after the release cells were infected with *C. pneumoniae* EBs (MOI 15) by centrifugation for 20 min at 37°C and fixed 1.5 h, 2 h or 2.5 h after release. The number of cells infected and the number of internalized EBs were determined via microscopy. **B** Determination of the number of *C. pneumoniae* infected cells at the indicated time points. Non-mitotic cells represent RO-3306-treated cells in interphase stages. Infected cells were defined as cells showing a minimum of one internalized EB. n=3, each representing 30 cells. Error bars denote ±SEM, p<0.01 (**), p<0.001 (***), p<0.0001 (****), ns; not significant (two-tailed unpaired Student’s *t*-test). **C** Quantification of the percentage of internalized *C. pneumoniae* EBs per cell. At least 16 cells with EBs were counted per time point and population. Error bars denote ±SEM, p<0.0001 (****), ns; not significant (two-tailed unpaired Student’s *t*-test). **D** Representative confocal fluorescence images of mitotic U2OS infected with *C. pneumoniae* EBs and fixed at the indicated time point after washout. MTs were visualized with anti-_α_-tubulin antibody (green), actin with rhodamine-phalloidin staining (red) and chromatin/EBs with DAPI (blue). White boxes show enlargements, white arrow heads point to internalized EBs. Scale bars: 10 µm. Each image consists of one layer of a z-stack. Beside images: Diagrammatic representation of the images shown: (i) actin ring (ii) chromatin (iii) spindle (vi) internalized EB.

DMSO mock-treated cells and RO-3306 treated cells were infected with *C. pneumoniae* EBs (MOI 15) 1 h after DMSO or RO-3306 washout. Cells were fixed 1.5 h, 2 h or 2.5 h after washout and analyzed by microscopy (experimental set-up shown in Fig 3A). EBs were able to infect mock-treated, mitotic and non-mitotic cells (Fig 3B). However, while 100% of the mock-treated cells and RO-3306 treated non- mitotic cells were infected by EBs, this was not the case for mitotic cells. At the first time point we analyzed (1.5 h after washout) only 18% of the mitotic cells were infected by *C. pneumoniae*; this number increased to 37% for the 2.5 h time point (Fig 3B). Furthermore, the number of internalized EBs in mitotic cells was significantly lower than in the other cell populations. The number of internalized EBs was nine and five-fold higher in non-mitotic and mock-treated cell populations versus mitotic cells at the 1,5 and 2,5 h time points, respectively (Fig 3C). Internalized EBs in mitotic cells were found in cells in various stages of mitosis with no specific localization inside the cell (Fig 3D, Fig S3C). Thus, the interphase MT cytoskeleton *per se* is an important contributor for chlamydial infection efficiency.

### *C. pneumoniae* infection leads to an increase of acetylated MTs

So far, we have found that the amount of acetylated interphase MTs in a cell correlated with early EB infection efficiency. This suggests a pre-permissive host cell state for pathogen entry. We next asked, if *C. pneumoniae* infection could alter host MT parameters. Thus, U2OS cells were infected with different MOIs ranging from 10 to 50 and the amount of acetylated MTs in such cells was determined. We used the GAPDH enzyme as an internal standard. 30 min pi the amount of acetylated MTs was similar in uninfected and infected cells irrespective of the MOI used (Fig 4A, quantification in 4D). In contrast, 1 hpi the amount of acetylated MTs in cells increased significantly and in a dose-dependent manner, while the total amount of MTs remained unchanged (Fig 4B, quantification in 4D, Fig S4). Infected cells had significantly more acetylated MTs than the non-infected cell population and the amount of acetylated MTs rose with increased MOI (Fig 4D). Thus, early infection by *C. pneumoniae* affects cellular MT composition. To test if the EBs actively caused this effect, we repeated the experiment using heat-inactivated EBs. A subset of heat-inactivated EBs can still bind to the host cell but fail to trigger the active infection processes [39, 41] (Fig S5A-C). No increase of acetylated MTs was observed when inactivated EBs were used irrespective of the MOI, suggesting that chlamydial heat-labile molecules are responsible for the increase of acetylated MTs (Fig 4C, D). Next, we tested if the internalization process of large particles such as *C. pneumoniae* EBs *per se* might be sufficient to increase the MT acetylation. Thus, we analysed MT acetylation in relation to internalisation of Invasin coated latex beads. Briefly, we used latex beads with a size of 1 µm coated with the *Yersinia pseudotuberculosis* effector protein Invasin tagged with His. Invasin is an adhesin that binds and activates multiple integrin-ß1 complexes including integrin α6β1 initiating internalization of the bacteria as well as that of latex beads coated with the protein [42, 43]. Using increasing amounts of coated beads that were taken up by the cell as shown by Western blot analysis, we found no difference in the amount of acetylated MTs (Fig 4E, F). We conclude that internalization of large particles *per se* is insufficient to trigger this response (Fig 4 E, F). Thus, early chlamydial infection results in increased MT acetylation suggesting that cytoskeletal MT remodelling is part of the initial establishment phase of infection.

**Fig 4.**
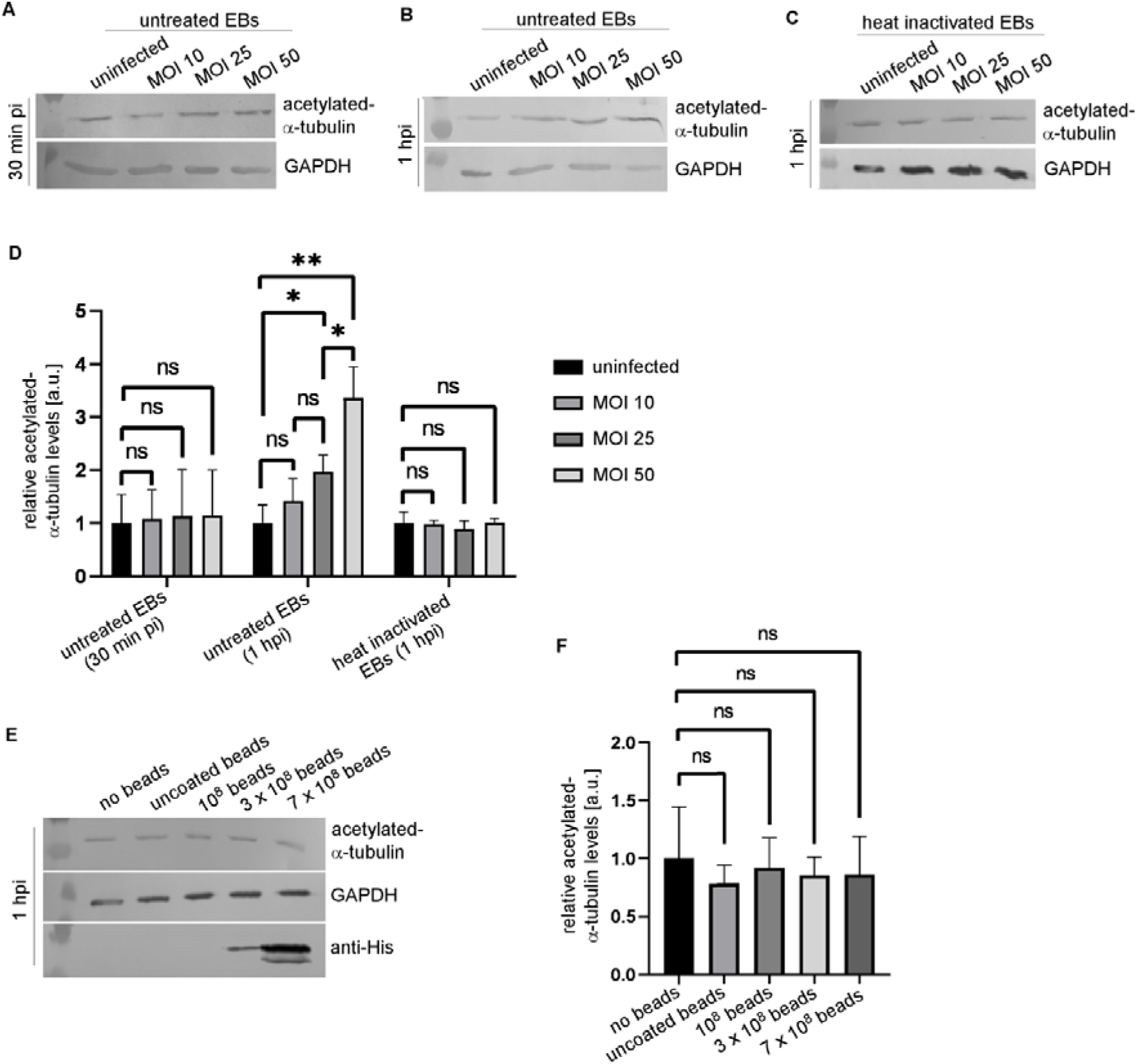
*C. pneumoniae* infection increases acetylated host MTs. **A** and **B** Western blot analysis of acetylated-_α_-tubulin in U2OS cells uninfected or infected with different MOIs of *C. pneumoniae* EBs. Cells were infected with *C. pneumoniae* EBs at the indicated MOI for 30 min (**A**) or 1h (**B**) followed by cell lysis. Protein extracts were separated on a 10% SDS-PAGE. Western blot analysis was carried out using either anti-acetylated-_α_-tubulin antibody or GAPDH antibody. The blots shown are representatives of three repeats/condition. **C** Western blot analysis of acetylated-_α_-tubulin in U2OS cells infected with the given MOIs using heat-inactivated EBs. Samples were treated as in **B**. The blot shown is a representative of three repeats. **D** Quantification of relative acetylated-_α_-tubulin levels shown in **A**-**C**. Band intensities from Western blots of acetylated-_α_-tubulin and GAPDH in cells infected with different MOIs of *C. pneumoniae* EBs were determined using ImageJ. The ratio of acetylated-_α_-tubulin levels to GAPDH levels was analysed and compared relative to control cells (The ratio for uninfected control set to 1). n=3 independent experiments/condition, Error bars denote ±SEM, p<0.05 (*), p<0.01 (**), ns; not significant (two-tailed unpaired Student’s *t*-test). **E** Western blot analysis of acetylated MTs in U2OS cells incubated with different numbers of Invasin coated latex beads. Cells were incubated with uncoated beads or the indicated number of beads coated with 20 µg recombinant His-tagged Invasin for 1h at 37°C before cell lysis. Protein extracts were separated on a 10% SDS-PAGE and Western blot analysis was performed using anti-acetylated-_α_-tubulin antibody, anti-GAPDH antibody or anti-His antibody. The blot shown is a representative of three repeats/condition. **F** Quantification of relative acetylated-_α_-tubulin levels shown in **E**. Band intensities from Western blots of acetylated-_α_-tubulin and GAPDH in cells incubated with latex beads were determined. The ratio of acetylated-_α_-tubulin levels to GAPDH levels was analysed and compared relative to control cells (The ratio for no beads set to 1). ImageJ was used for band intensity quantification. n=3 independent experiments/condition, Error bars denote ±SEM, ns; not significant (two-tailed unpaired Student’s *t*-test).

### The early secreted effector protein CPn0572 suppresses force-dependent MT catastrophe

The findings that host cell MT acetylation is increased by *C. pneumoniae* infection points to a possible involvement of early chlamydial effector proteins in this process. As we previously identified a large number of *C. pneumoniae* proteins that alter the MT cytoskeleton among them nine proteins coded for by early and late expressed genes, it is likely that several effector proteins jointly manipulate the MT cytoskeleton in the early infection [27]. To start to understand how MTs might be modulated by chlamydial proteins at this very early infection stage, we assayed how CPn0572 an early effector protein of the TarP family alters MT cytoskeleton parameters. The chlamydial TarP family are major early effectors that physically associate with and remodel the host actin cytoskeleton [7, 44] with CPn0572 having a unique additional function in also binding to and modulating the MT cytoskeleton [29]. In fact, ectopic expression of a CPn0572 variant that contains the MT binding region, resulted in an increase of MT acetylation in HEp-2 cells [29]. To determine, how CPn0572 might affect MT dynamics in living cells, we expressed a genome-integrated *cpn0572-mCherry* version via the inducible TetO promoter system in a fission yeast *Schizosaccharomyces pombe* strain which allowed live-cell imaging of the MT cytoskeleton [45, 46] (Fig 5A). The reason for this experimental set-up was as follows: (i) we used single-locus TetO-driven genomic expression to minimize dosage heterogeneity and capture early effects of CPn0572 on interphase MT dynamics and importantly (ii) benefited from the well understood, simple *S. pombe* MT cytoskeleton, where MTs are organized in discrete MT bundles whose dynamic behaviour can be measured easily using life-cell imaging of α-tubulin-GFP [47, 48]. We have previously shown that chlamydial MT modulating proteins can be identified and studied in *S. pombe* [27, 28]. The 3-4 *S. pombe* interphase MT bundles consist of MT filaments with MT plus-ends oriented toward the cell tips and MT minus-ends at the MT organizing centres near the nucleus. Two antiparallel bundles overlap at the cell centre. MT plus-ends grow from the cell centre along the long axis to the cell tip, where MT polymerization stops, followed by pausing, a MT catastrophe event and depolymerization (Fig 5B). This cycle is then repeated numerous times. MT dynamics of α-tubulin-GFP cells that were between 9-10 µm cell length and expressing CPn0572-mCherry via TetO for 1 h (Fig 5C, bottom) were compared to α-tubulin-GFP expressing wild-type cells of the same length (Fig 5C, top) [46]. At this time point, cells expressing CPn0572 remained fully viable and 93% of these CPn0572-mCherry expressing cells showed a highly similar pattern: one to three discrete CPn0572-mCherry MT-associated foci (Fig 5C and Fig S6A-D). Localization of these foci was not uniform but the majority was found at MT overlap zones. MT polymerization rate in these cells was not altered in comparison to the wild-type control strain (Fig 5D), but the rate of MT depolymerization and the number of catastrophe events were reduced significantly (Fig 5E, F). Additionally, we found that in CPn0572 expressing cells MT organization was altered: the MT cytoskeleton was more disorganized (Fig 5G, Fig S7A) with the wild-type MT spatial distribution along the long axis of the cells reduced (Fig 5H, Fig S7B).

**Fig 5.**
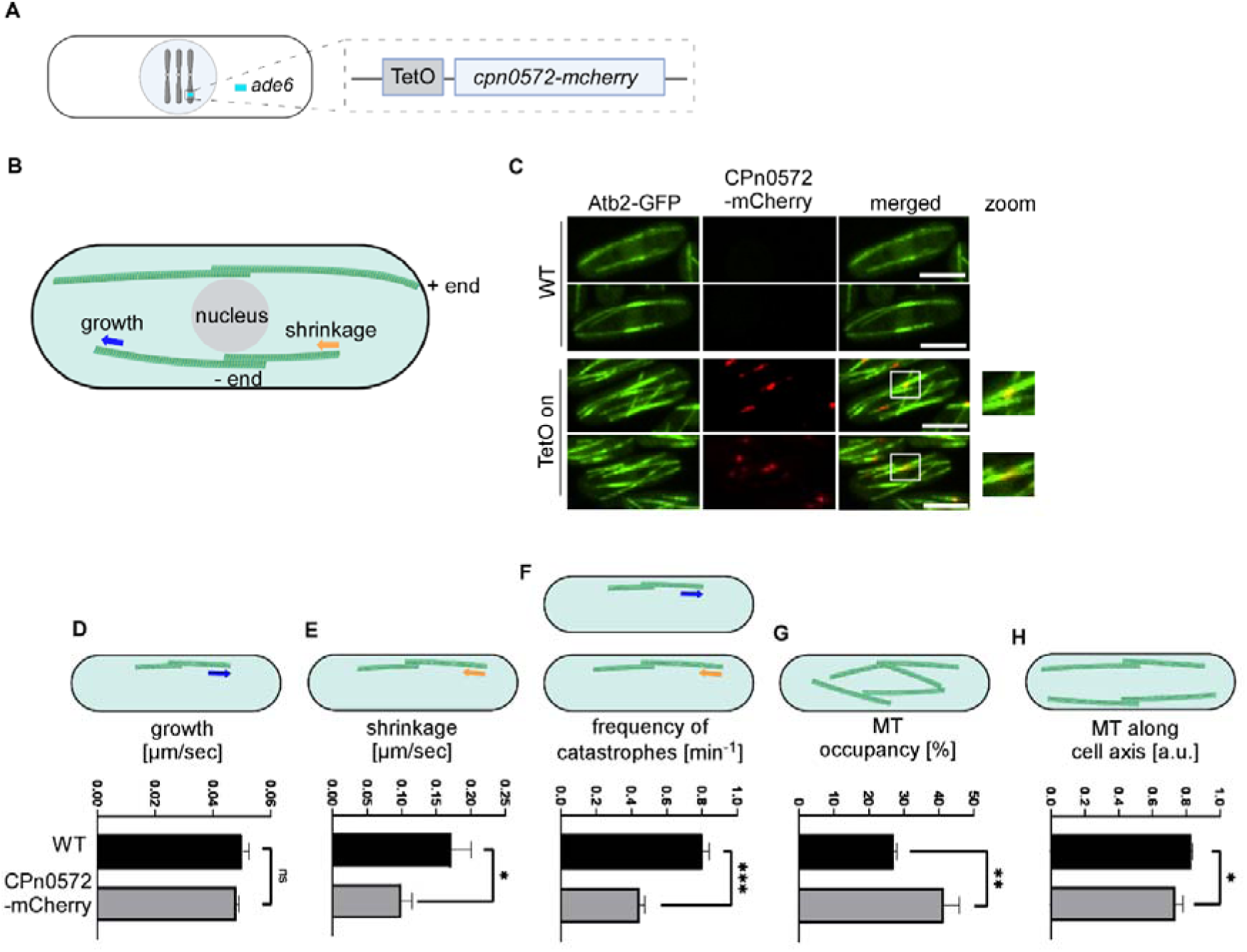
CPn0572 alters *S. pombe* MT dynamics and organisation. **A** Schematic representation of genome-integrated *cpn0572-mcherry* under control of the TetO promotor integrated in the *ade6^+^* locus of *S. pombe*. **B** Diagrammatic representation of *S. pombe* MT bundle behaviour. Explanation in main text. Blue arrow: polymerizing bundle; orange arrow: depolymerizing bundle. **C** Confocal live-cell images of wild-type α-tubulin-GFP (Atb2-GFP) and CPn0572-mCherry α-tubulin-GFP expressing cells. Top, wild-type cells; Bottom, CPn0572-mCherry expressing cells. Expression was induced by addition of 2,5 µg tetracycline for 1 h to logarithmically growing cells at 30°C. White boxes show enlargement of merged images. Scale bar: 5 µm. **D**-**H** Quantification of interphase MT dynamics and organization in wild-type and CPn0572-mCherry expressing cells. MT growth and shrinkage rates (n=3 biological replicates with at least 16 MTs) and the frequency of catastrophes (n=3 biological replicates with at least 16 MTs) were quantified from time-lapse images. MT occupancy i.e. spatial organisation within cell was determined as shown in Fig S7A. MT organisation along the long axis of the cell was determined as shown in Fig S7B. Data were quantified from images taken at 22 °C using ImageJ (n=3 biological replicates with at least 28 cells). Error bars denote ±SEM, p<0.05 (*), p<0.01 (**), p<0.001 (***), ns; not significant (two-tailed unpaired Student’s *t*-test).

Importantly, we found that CPn0572 suppresses cell tip-induced catastrophe. Interphase MT plus-ends in wild-type cells polymerize to the cell end and upon contact with the cortex will pause followed by a catastrophe event and depolymerization (Fig 6A, B, top panels). Individual MT bundle behaviour of wild-type MT bundles at the cell tip is shown in Fig 6C. However, in CPn0572-mCherry expressing cells, the normal local regulation at the cell tip was suppressed and instead such MT bundles continued to polymerize “curling” around the cell tip (Fig 6A, B, bottom panels). Behaviour of individual bundles in a CPn0572-expressing strain is depicted in Fig 6D and comparison of typical wild-type and CPn0572 bundles behaviour is shown in Fig 6E. Quantification of MT dynamics in wild-type and CPn0572-expressing cells revealed that the dwell time at the cell end was significantly increased in the latter cells, while the time of pausing was significantly reduced (Fig 6F, G).

**Fig 6.**
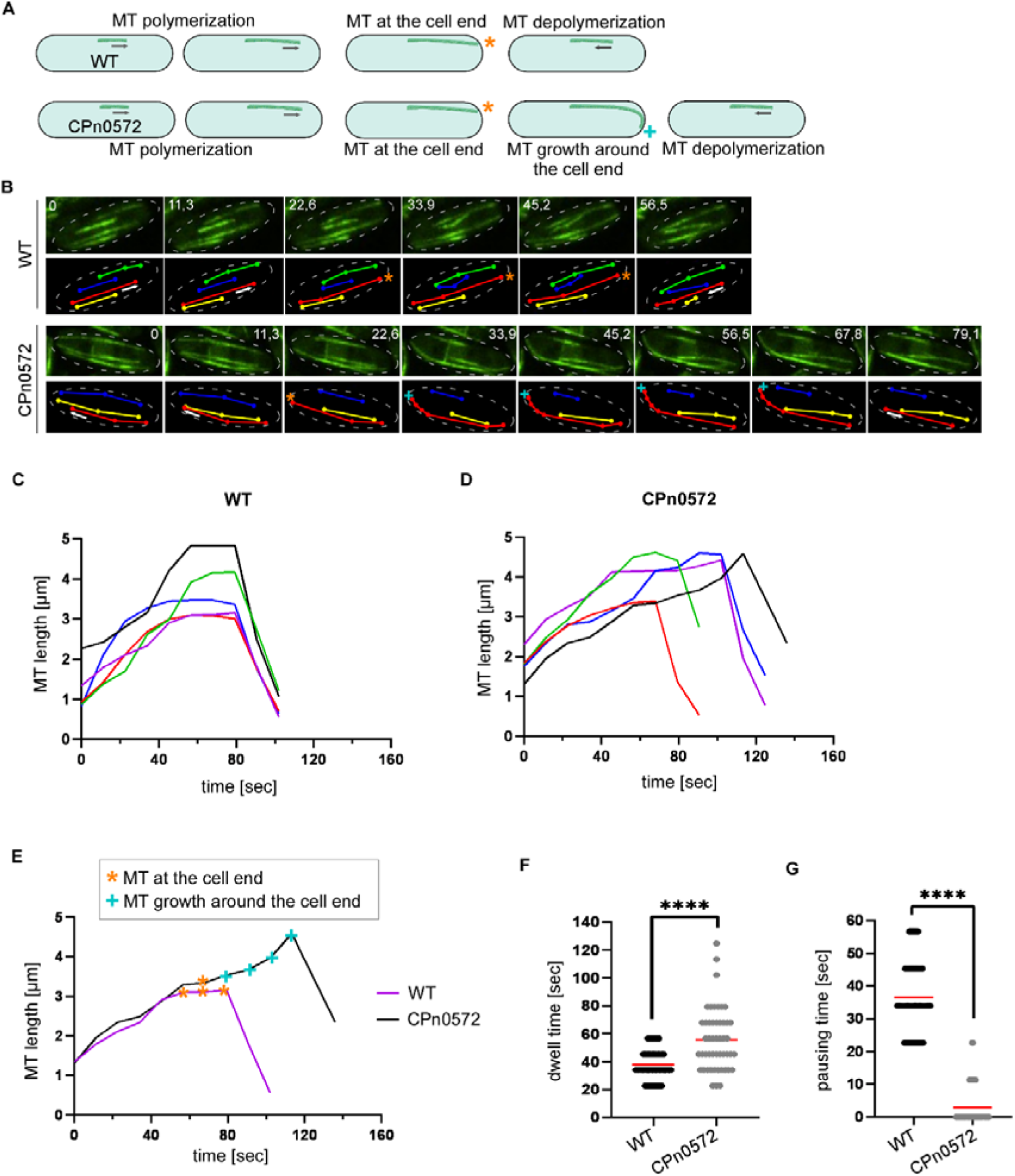
CPn0572 promotes MT persistence at a MT catastrophe site. **A** Schematic representation showing interphase MT dynamics of the indicated *S. pombe* strains. Top panels: Wild type MT bundles polymerize along the long axis of the cell toward the cell tip. Reaching the cell end, MTs pause (orange asterix) followed by a catastrophe event and then depolymerize. MT +ends (plus-ends) of CPn0572-mCherry expressing cells do not pause at the cell tip but continue growing around the cell end (cyan cross) before a catastrophe event followed by depolymerization. **B** Time-lapse confocal images and the corresponding MT tracks to show MT behaviour of the indicated *S. pombe* strains. CPn0572-mCherry expression was induced 1 h before starting microscopy. Images were taken every 11,3 seconds and are shown as maximum intensity projection. Images shown for WT are generated from movie 1 and images for CPn0572 from movie 2 in the supplement. **C** and **D** MT dynamics of individual MT bundles from wild-type (WT) or CPn0572-mCherry expressing cells. Each colour graph shows represents an individual MT bundle from an individual cell. MT bundles chose for analysis showed a length of at least ∼1 µm at the beginning of measurement. **E** Representative examples of the MT behaviour of wild-type (WT) and CPn0572-mCherry expressing cells. **F** Quantification of the dwell time of wild-type cells and CPn0572-mCherry expressing cells. The dwell time represents the time a MT bundle remains at the cell end including pausing and continued growth. 50 individual MTs were analysed/strain. The red bar represents the mean. p<0.0001 (****) (two-tailed unpaired Student’s *t*-test). **G** Quantification of the pausing time of 50 individual MTs in wild-type and CPn0572-mCherry expressing cells. The pausing time represents the duration a MT bundle stays at the cell end without polymerizing/depolymerizing. The red bar represents the mean. p<0.0001 (****) (two-tailed unpaired Student’s *t*-test).

The combined reduction in catastrophe frequency and depolymerization rate, together with continued growth at cell ends, suggests that CPn0572 increases MT stability and persistence, particularly under conditions that normally trigger catastrophe. Thus, we propose that early chlamydial effector proteins can locally alter MT behaviour to impact the infection process.

## Discussion

### Pre-existing microtubule architecture promotes efficient *C. pneumoniae* entry

Our data identify a previously unrecognized relationship between the pre-existing MT state of the host cell and the efficiency of *C. pneumoniae* EB entry. In a cold-recovery assay, EBs preferentially infected cells containing acetylated MTs, and this effect increased in a dose-dependent manner. In the same assay, different amounts of detyrosinated MTs did not have this effect. Thus, we propose that productive EB entry is favoured by a specific pre-existing MT architecture of the host cell. To our knowledge, the present study provides the first evidence that the MT subpopulation of a cell present before or at the moment of infection influences bacterial entry or very early infection events occurring within minutes of pathogen-host cell contact.

Importantly, our pharmacological experiments show that tubulin acetylation itself is unlikely to be the determining factor. While stabilization of MTs via Taxol increased the number of acetylated MTs and EB entry, increasing tubulin acetylation through HDAC6 inhibition with Tubacin did not. Taxol stabilisation of MTs leading to MT acetylation may not fully recapitulate the properties of naturally acetylated MTs, since physiological acetylation occurs within the broader context of the tubulin code and associated regulatory factors [33, 34, 36, 49]. However, EB internalisation of cells from the cold-recovery assay and EB internalisation of Taxol-treated cells again show dose-dependency. Taxol-treated cells have more acetylated MTs than cells from the cold-recovery and concurrently the highest number of internalized EBs.

Due to their increased lifetime and resistance to mechanical forces, acetylated MTs are known to serve as tracks for vesicular traffic e.g. within cells of the innate immune system [50]. During phagocytic events, such as the chlamydial EB entry, membrane-containing vesicles are transported to the site of phagosome formation, thereby providing sufficient endomembrane material for the engulfment and internalization of the pathogen [51]. Such MT-dependent supply could provide a local increase of membrane material resulting in more efficient EB uptake. Indeed, Chlamydia infect epithelial cells via clathrin-dependent processes [52]. Clathrin-mediated receptor endocytosis typically generates vesicles with a diameter of 120 nm [53], thus chlamydial EBs with a diameter of 400 nm require a significantly larger amount of membrane for their formation. In fact, during the internalization of *C. pneumoniae* EBs, invading EBs come into contact with EGFR-positive structures as early as 5 min pi [39]. 1 hpi the juvenile inclusion formed around the EB is loaded with EGFR and often EGFR-positive vesicles are associated with the early inclusion or in the process of fusion with it [39]. As a result, the juvenile inclusion membrane typically lies very loosely around the internalized EB [39]. In addition, MTs play an important role in regulating signalling pathways, including those responsible for actin organization. In a variety of cell types including epithelial cells, MT depolymerization leads to the release of GEF-H1 which in turn activates RhoA the antagonist of Rac1 [54, 55]. Thus, stable MTs might alter the balance away from the GEF-H1/RhoA axis, hereby indirectly supporting Rac1-induced actin structures required for protrusion formation, necessary for EB internalization.

### Chlamydia remodels the host microtubule cytoskeleton during early infection

The preferential infection of cells containing acetylated MTs needs to be distinguished from the increase in MT acetylation observed 1 h after chlamydial infection. The former identifies a host-cell property that exists prior to infection, the latter observation demonstrates that *C. pneumoniae* alters the host MT cytoskeleton during early infection. Infection increased acetylated α-tubulin in a dose-dependent manner, consistent with enhanced MT stability.

What selective advantage would an intracellular pathogen gain from creating a more long-lived, acetylated MT network immediately after entry? For obligate intracellular pathogens such as *Chlamydia pneumoniae*, efficient intracellular trafficking is essential immediately following host cell entry. Newly internalized Chlamydia-containing vacuoles termed inclusions originate at the cell plasma membrane and subsequently undergo dynein-dependent transport along MTs toward the centrosomal/perinuclear region, where the replication niche is established [11]. Thus, successful infection depends on access to a functional MT network capable of supporting long-range motor-driven transport. This is all the more important given that the chlamydial EB containing vesicle is much larger than a typical early endosome and small endosomal vesicles move faster than larger ones [53, 56]. In this context, the infection-induced increase in acetylated α-tubulin observed during the first hour of infection may be biologically significant. Acetylated MTs are enriched within long-lived MT populations that support long-range intracellular transport. Furthermore, α-tubulin K40 acetylation increases MT resilience to repeated mechanical stress, thereby protecting MTs from mechanical ageing and lattice breakage [33]. As a result, acetylated MT can maintain persistent intracellular transport routes over extended periods. Although it is not known whether dynein directly recognizes acetylated MTs, the generation of a long-lived and mechanically resilient MT network would result in efficient motor-dependent trafficking. We therefore propose that early effector-mediated alteration of the host MT cytoskeleton may facilitate transport of chlamydial inclusions toward the centrosomal region required for the establishment of the intracellular replication niche.

The rapid and dose-dependent increase in acetylated MTs observed within 1 hpi suggests that *C. pneumoniae* EBs deliver preformed effector proteins capable of modulating the host MT cytoskeleton immediately upon host cell contact. To date no effector proteins have been pinpointed for such a task, but *C. pneumoniae* CPn0572 and CPn0216 represent two strong candidates. CPn0572, a member of the chlamydial TarP family is secreted via the type III secretion system and can therefore access the host cytoplasm during the earliest stages of infection [7]. CPn0572 directly targets the host cytoskeleton by associating *in vivo* and *in vitro* directly with both actin filaments and MTs. In mammalian cells, ectopic expression of CPn0572 induces prominent MT bundles, while during infection secreted CPn0572 localizes to MTs. Importantly, ectopic expression of a CPn0572 variant that exclusively binds MTs resulted in hyper-stabilized MTs and an increase in acetylated MTs [29]. The second candidate is CPn0216, which is unique to *C. pneumoniae.* We identified this protein in a systematic *S. pombe* screen as one of 13 chlamydial proteins that modulate the MT cytoskeleton; however, CPn0216 was unique in its ability to promote MT stabilization [27]. Although CPn0216 is classified as an Inc protein localizing to the inclusion membrane during infection, its late expression during the chlamydial developmental cycle raises the possibility that it accumulates in EBs and is therefore already present at the onset of infection [57]. The ability of both CPn0572 and CPn0216 to stabilize MTs makes them plausible contributors to the rapid increase in MT acetylation observed shortly after EB uptake.

However, it is likely that the observed dose-dependence increase in MT acetylation observed 1 hpi is caused by multiple EB-associated effectors that together remodel the host MT network. Importantly, the *S. pombe* screen was inherently limited to chlamydial proteins whose ectopic expression was tolerated by the yeast cell. Indeed, CPn0572 was not included in the screen because its expression is detrimental to fission yeast cells [28]. Thus, the MT-modulating proteins identified to date [27] likely represent only a subset of the MT-targeting repertoire encoded by *C. pneumoniae* and further MT-modulating *C. pneumoniae* effectors exist.

### CPn0572 alters microtubule boundary-response behaviour

CPn0572 is one of the candidate chlamydial effectors that have a MT stabilizing effect. It was thus interesting to analyse how CPn0572 alters MT dynamics via life-cell imaging. In particular, the use of the inducible, genome-integrated TetO promoter to express CPn0572 was important. Although TetO-driven expression cannot recapitulate the spatially restricted injection of an effector by a bacterial secretion system, it allows controlled temporal induction of CPn0572-mCherry in live *S. pombe* cells. Thus, it provides a tractable approximation of effector appearance after secretion, enabling analysis of downstream effects on MT dynamics. TetO-controlled CPn0572-mCherry was present on MTs as foci in close proximity to MT overlap zones and its presence altered MT dynamics and spatial distribution. Most notably, MTs reaching the cell tip failed to undergo the characteristic catastrophe response associated with cortical contact and instead continue to grow along the cell boundary. In *S. pombe*, interphase MTs plus-ends grow towards the cell ends, where cortical contact and compressive forces contribute to catastrophe induction [48, 58, 59]. This response is also regulated by MT-associated proteins. For example, the kinesin-8-member Klp5 promotes catastrophe at cell ends and absence of this protein results in an MT cell end phenotype similar to that seen in CPn0572 expressing cells [60]. Thus, the continued growth of MT plus-ends around the cell tip in CPn0572 expressing cells, suggests that CPn0572 interferes with the normal boundary-induced catastrophe response.

Importantly, CPn0572-mCherry signals are not enriched at MT plus-ends but localized preferentially to MT overlap regions. This suggests that CPn0572 affects catastrophe indirectly, possibly by altering MT bundle architecture, lattice stability, or force transmission within antiparallel bundles. Thus, we suggest that CPn0572 acts as a cytoskeletal organizer rather than a canonical MT plus-end regulator. During early infection, such activity could contribute to the coordinated remodelling of host actin and MTs during bacterial uptake. We propose that CPn0572 promotes a MT state that is less prone to boundary-induced catastrophe, thereby favouring persistent MT interactions with membrane- or actin-associated structures encountered during the earliest stages of chlamydial infection. Since CPn0572 modulates both actin and microtubules, it is tempting to speculate that the increased MT persistence observed in CPn0572-expressing cells promotes prolonged interactions between MTs and actin-rich structures. Actin–MT crosstalk is frequently associated with the generation of stabilized MT populations in polarized cells [61], raising the possibility that CPn0572 contributes to the accumulation of acetylated MT observed during chlamydial infection.

## Material and Methods

### Cell culture of mammalian cells and bacterial strains

Human epithelia U2OS cells (ATCC; HTB-96) were cultured in Dulbecco’s modified Eagle’s medium (DMEM; Thermo Fisher Scientific) supplemented with 10% fetal calf serum (FCS), MEM vitamins (Thermo Fisher Scientific) and non-essential amino acids (Thermo Fisher Scientific) at 37°C and 6% CO2. *C. pneumoniae* GiD [62] was propagated in U2OS cells. EBs were purified in 30% gastrographin solution (Bayer Vital GmbH, Leverkusen, Germany) and stored at -80°C in SPG buffer (3.8 mM KH_2_PO_4_, 10.8 mM Na_2_HPO_4_, 4.9 mM L-glutamine, 220 mM sucrose).

### *C. pneumoniae* infection of mammalian cells

Human epithelial U2OS cells were cultivated in supplemented DMEM medium and incubated at 37°C and 6% CO_2_. 50% confluent cells on coverslips in 24 well plates were covered with supplemented DMEM medium including *C. pneumoniae* EBs at a given multiplicity of infection (MOI) and centrifuged at 2,980 rpm for 60 min at 37°C. The medium was replaced with fresh medium containing 12 µg/ml cycloheximide and incubated at 37°C and 6% CO_2_. Paraformaldehyde-fixed samples were used for microscopy. After permeabilization with 2% saponin solution, chlamydial inclusions were visualized with *C. pneumoniae* Inc protein antibody anti-CPn0147 (generated in our lab, 1:50) and DNA was stained with DAPI (Merck KGaA, Darmstadt, Germany, 1:500). For early infection (< 2h) 70% confluent cells in 6 well plates or on coverslips in 24 well plates were infected with *C. pneumoniae* EBs at the given multiplicity of infection (MOI) in supplemented DMEM medium and centrifuged at 2,980 rpm for 20 min at 37°C or 4°C. Cells were then incubated without media change at 37°C and 6% CO_2_. Cells in 6 wells were washed three times with HBSS (Thermo Fisher Scientific), lysed with Phospho-Lysis buffer (1% NP40, 1% Triton X100, 20 mM Tris, 140DmM NaCl, 2 mM EDTA, 1DmM Na_2_VO_4_, Roche Protease Inhibitor Cocktail) and analysed by Western blot. 24 well samples were fixed with 3% paraformaldehyde and prepared for antibody staining.

### Western blot analysis

Whole cell lysates from U2OS cells were prepared with 4xSDS Buffer (200 mM Tris pH 6.9, 8% SDS, 0.2% Bromphenol blue, 20% Glycerin) and DTT (Merck KGaA, Darmstadt, Germany, 100 µM). *S. pombe* cells were lysed in lysis buffer (1xSDS Buffer, 100 mM DTT) resulting in whole cell lysates ready to use for western blot analysis. Equal volumes (20 µl) were loaded on a 10% SDS-PAGE and transferred to PVDF membranes using a semi-dry blotting system. Blocking was performed in blocking solution (5% milk in PBS, 0.05% Tween) for 1h at RT. Western blotting was carried out using the following primary antibodies: anti-α-tubulin (Mouse, Merck KGaA, Darmstadt, Germany, 1:1000), anti-acetylated-α-tubulin (Mouse, Thermo Fisher Scientific, 1:20000), anti-GAPDH (Mouse, Merck KGaA, Darmstadt, Germany, 1:1000), anti-His (Mouse, Santa Cruz Biotechnology, Inc., 1:200) and anti-mCherry (Mouse, Thermo Fisher Scientific, 1:1000). Secondary antibody was AP conjugated anti-mouse IgG (Goat, Thermo Fisher Scientific, 1:30000).

### Cold- and/or drug-induced MT alteration

For cold-induced MT depolymerization, U2OS cells were placed on ice for 30 min und MT status determined microscopically using fixed cells. After ice treatment, cells were incubated at 37°C for 0-20 min followed by fixation using 3% paraformaldehyde to determine MT repolymerization. MTs were stained with anti-α-tubulin antibody (Merck KGaA, Darmstadt, Germany, 1:1000), acetylated MTs with anti-acetylated-α-tubulin antibody (Thermo Fisher Scientific, 1:200), detyrosinated MTs with anti-detyrosinated-tubulin antibody (Thermo Fisher Scientific, 1:200) and DNA was visualized with DAPI (Merck KGaA, Darmstadt, Germany, 1:500). AlexaFlour488 (Thermo Fisher Scientific, 1:2000) was used as secondary antibody. For very early infection studies, cells were infected with *C. pneumoniae* EBs with MOI 10 after ice treatment by centrifugation at 4°C and then incubated for 10 min at 37°C before 3% paraformaldehyde fixation.

For drug-induced MT alteration, U2OS cells were incubated with 10 µM Taxol (Merck KGaA, Darmstadt, Germany) or 10 µM Tubacin (Enzo Biochem Inc., New York) for 2h at 37°C and MT acetylation determined via Western blot analysis. Drug-treated cells were placed on ice for 30 min, fixed with 3% paraformaldehyde and MTs were visualized microscopically using an anti-α-tubulin antibody (Merck KGaA, Darmstadt, Germany), acetylated MTs with an anti-acetylated-α-tubulin antibody (Thermo Fisher Scientific) and DAPI (Merck KGaA, Darmstadt, Germany) was used to stain DNA. U2OS cells incubated with 10 µM Taxol (Merck KGaA, Darmstadt, Germany) or 10 µM Tubacin (Enzo Biochem Inc., New York) for 2h at 37°C were infected with *C. pneumoniae* EBs at 37°C by 20 min centrifugation at 37°C. Cells were fixed with 4% paraformaldehyde and internalized EBs were quantified via inside-out staining.

### Inside-out staining

Drug treated cells were infected with *C. pneumoniae* EBs for 10 min after centrifugation at 37°C as described above. To stain non-internalized EBs, paraformaldehyde-fixed cells were incubated with an anti-GiD antibody (generated in our lab, 1:40) in 1xPBS (137 mM NaCl, 2.7 mM KCl, 10 mM Na_2_HPO_4_, 1,8 mM KH_2_PO_4_) for 30 min at 30°C and then washed three times with 1ml 1xPBS. For secondary antibody AlexaFluor594 (Thermo Fisher Scientific, 1:1000) in 1xPBS was added to the cells, which were then incubated 30 min at 30°C followed by washing 3 times with 1xPBS. To stain internalized EBs, these cells were permeabilized with 2% saponin solution for 20 min at RT and then incubated with anti-GiD antibody (generated in our lab, 1:40) in 0,5% saponin solution for 30 min at 30°C. After three washing steps with 1 ml 0,5% saponin solution cells were incubated with AlexaFluor488 antibody (Thermo Fisher Scientific, 1:2000) in 0,5% saponin solution for 30 min at 30°C and again washed three times with 1 ml 0,5% saponin solution. Non-internalized EBs are now shown in yellow, all internalized EBs are visualized in green.

### Arrest of mammalian cells at G_2_/M boundary due to CDK1 inhibition

70% confluent U2OS cells, cultivated in 24 well plates on coverslips in supplemented DMEM medium, were incubated at 37°C with 6 µM CDK1-inhibitor RO-3306 (Merck KGaA, Darmstadt, Germany) for 20 h to block cells at G_2_/M. For drug-removal cells were washed twice with supplemented DMEM medium and incubated at 37°C with fresh medium. 1 h after drug washout cells were infected with *C. pneumoniae* EBs (MOI 10) by centrifugation for 20 min at 37°C and incubated at 37°C and 6% CO_2_ for 1,5 h to 2,5 h. Cells were fixed with 3% paraformaldehyde solution and prepared for microscopy. After permeabilization with 2% saponin solution rhodamine-phalloidin (Thermo Fischer Scientific, Waltham, MA, USA; #R415; 0,5 µl of 400X stock solution in 200 µl of PBS-Saponin solution for each coverslip) was used to visualize actin, anti-α-tubulin antibody for MTs (Merck KGaA, Darmstadt, Germany) and DAPI for DNA/EBs (Merck KGaA, Darmstadt, Germany). For secondary antibody AlexaFluor488 (Thermo Fisher Scientific, 1:2000)

### Yeast strains and growth conditions

The following *S. pombe* strains were used in this study:

#605: *h^-^*, *his3-D1*, *ade6-M210*, *leu1-32*, *ura4-D18*

#2334: *h^+^*, *leu1-32::SV40::atb2^+^-GFP[LEU2]*, *ura4-D18*, *his3-D1*

#3779: *h^-^*, *ade6^+^::P.CMV-tetR:PenotetS-CPn0572-mCherry-Scer\T.ADH1:hphMX*, *his3-D1*, *leu1-32*, *ura4-D18*

#3799: *ade6^+^::P.CMV-tetR:PenotetS-CPn0572-mCherry-Scer\T.ADH1:hphMX*, *leu1-32::SV40::atb2^+^-GFP[LEU2]*, *his3-D1*, *ura4-D18*

*S. pombe* strains were cultured at 30°C in rich medium (YE5S) or minimal medium (plus appropriate supplements) [63]. For microscopy, *S. pombe* strains were grown in liquid minimal medium at 30°C to mid-log phase. Expression of CPn0572-mCherry was induced by adding 2,5 µg Anhydrotetracycline (Thermo Fisher Scientific) 0-3 h before microscopy.

To generate the CPn0572-mCherry expressing strain (#3779), *cpn0572-mcherry* fusion was amplified from a plasmid and the DNA fragment cloned between the BglII and NheI sites in the integration plasmid pDB5318 replacing mECitrine (Addgene plasmid #204828) [45]. The resulting plasmid was named p1713. For genome integration of *cpn0572-mcherry* p1713 was linearized with NotI and transformed into strain #605 where it integrated into the *ade6^+^* locus via homologous recombination (selection via Hygromycin B resistance). Colony PCR was used to verify the correct integration. The strain was backcrossed once and received the number 3779. To generate a strain expressing CPn0572 and α-tubulin-GFP, strains #2334 and #3779 were mated and the resulting spores tested for the presence of *leu1-32::SV40::atb2^+^-GFP[LEU2]* (leucine prototrophy) and *ade6^+^::P.CMV-tetR:PenotetS-CPn0572- mCherry-Scer\T.ADH1:hphMX* (Hygromycin B resistance)(Hygromycin from Merck KGaA, Darmstadt, Germany, 100 µg/ml).

### Microscopy and image processing

For mammalian cells, microscopic images were acquired using an inverse Nikon TiE Live Cell Confocal C2plus with 100-x TIRF objective and a C2 SH C2 Scanner. For live cell imaging of *S. pombe*, cells were mounted on agar pads as described previously [64]. Images were acquired by using a confocal Evident SpinSR microscope with 100-x oil immersion objective (Olympus, Tokyo, Japan). Image editing was done with ImageJ 1.47v (National Institute of Health).

### Data analysis

Image analysis and Western blot quantification were done using ImageJ 1.47v (National Institute of Health, USA). For colocalization analysis of MT PTM versus all MTs the Coloc2 Plugin of ImageJ was used. The protocol for Western blot quantification can be found here: https://www.yorku.ca/yisheng/Internal/Protocols/ImageJ.pdf

Data analysis and statistical significance were performed with Prism (GraphPad Software Inc., USA).

## Acknowledgement

We thank Dr. Roland Piekorz for the U2OS cell line, Dr. Sebastian Hänsch at the Center for Advanced Imaging (CAi) for his help with microscopy and Dr. Abel Alcázar-Román for discussing experimental procedures (all at Heinrich-Heine-University, Düsseldorf, Germany). This study was supported by the Jürgen Manchot Stiftung, Düsseldorf, Germany to KS, http://www.manchot.org. The funders had no role in study design, data collection and analysis, decision to publish or preparation of the manuscript.

