## Supplementary material for "A Distinct Interphase Microtubule State Marks Host Cell Permissiveness to *Chlamydia pneumoniae* Entry"

Supplementary figures and legends

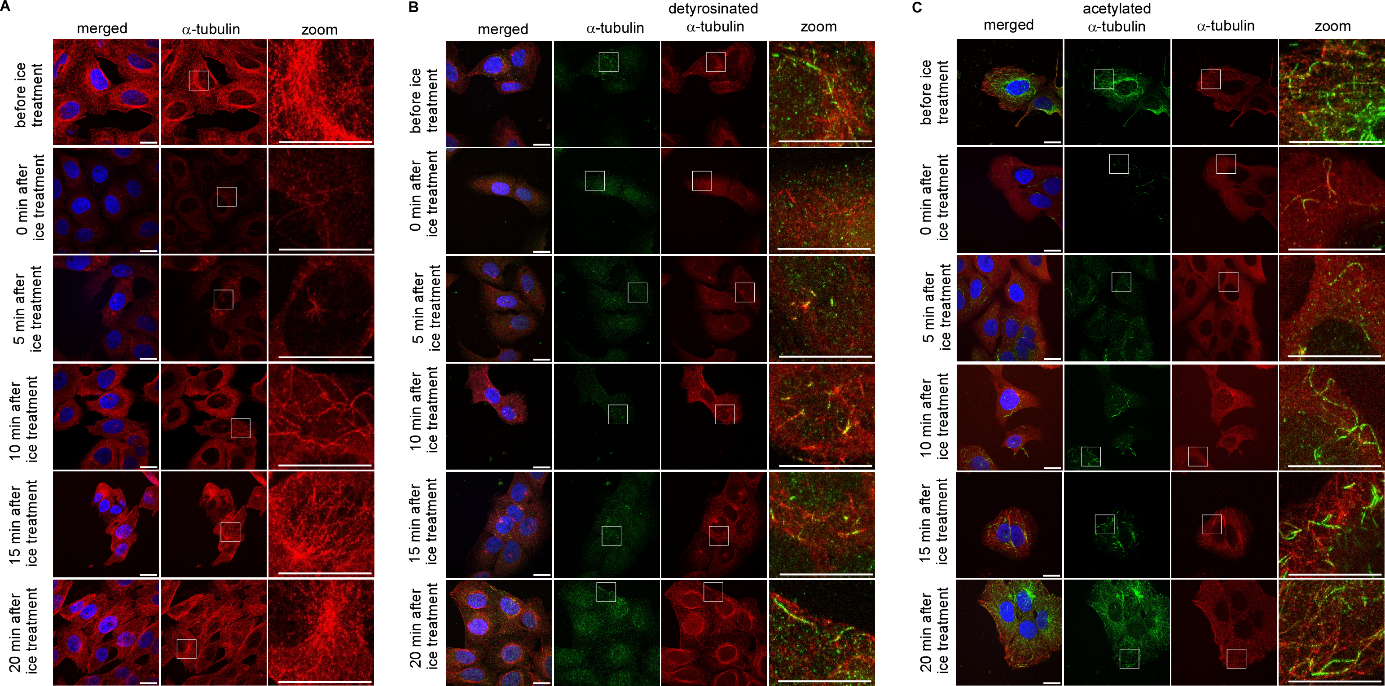

**Fig S1. MT phenotypes after ice treatment**

**A** Representative confocal fluorescence images of MTs in U2OS cells before, at and after ice treatment. Cells were incubated on ice for 30 min to depolymerize MTs followed by 0-20 min incubation at 37°C to allow MT repolymerization. MTs were visualized with an anti-α-tubulin antibody (green) and DNA is stained with DAPI (blue). Scale bars: 10 µm. **B** Representative confocal fluorescence images of detyrosinated MTs in U2OS cells before and after ice treatment. Cells were treated as described in **A**. MTs are shown in red with an anti-α-tubulin antibody, detyrosinated MTs in green with an anti-detyrosinated-tubulin antibody and DNA is visualized in blue with DAPI. Scale bars: 10 µM. **C** Representative confocal fluorescence images of all MTs and acetylated MTs in U2OS cells before and after ice treatment. Cells were treated as described in **A**. MTs stained with an anti-α-tubulin antibody (red); acetylated MTs with an anti-acetylated-α-tubulin antibody (green) and DNA with DAPI (blue). Scale bars: 10 µM. **A**-**C** Images are shown as maximum intensity projection.

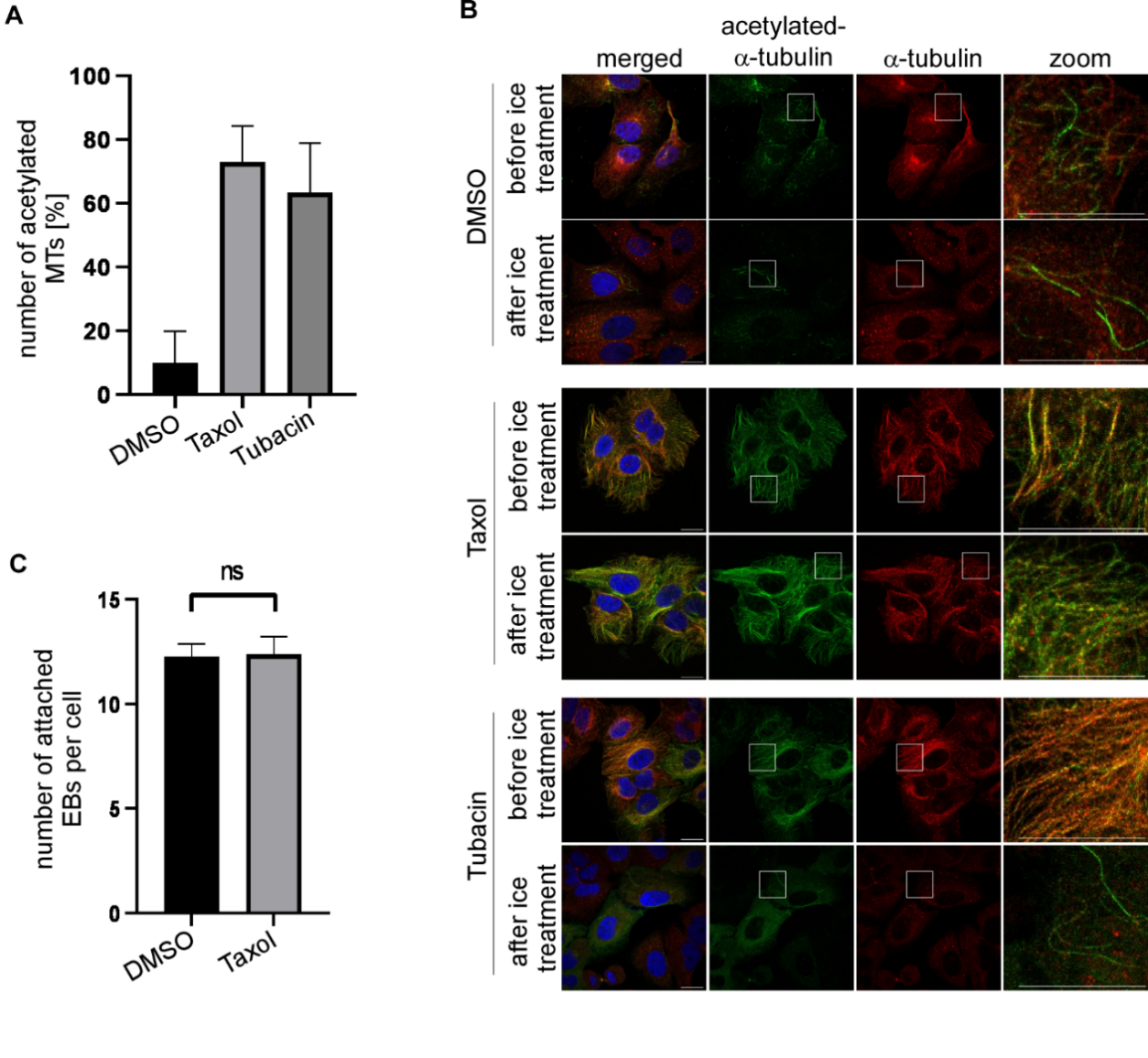

**Fig S2. Drug-induced MT alteration and EB adhesion**

**A** Quantification of the number of acetylated MTs in cells. The ImageJ Coloc2 Plugin was used to determine the amount of the acetylated MTs/all MTs as descripted in images similar to those shown in Fig 1B. n=15 cell per condition, Error bars denote ±SEM. **B** Representative confocal images of U2OS cells treated with DMSO (0,1%), Taxol (10 µM) or Tubacin (10 µM). Cells were incubated with the specific drug or DMSO for two hrs and fixed (labelled before ice-treatment) or incubated on ice for 30 minutes after drug/DMSO-treatment and then fixed (labelled after ice- treatment). All MTs are visualized with an anti-α-tubulin antibody (red), acetylated MTs with an anti-acetylated-α-tubulin antibody (green) and DNA is stained with DAPI (blue). Scale bars: 10 µM. **C** Quantification of the number of attached EBs per cell. EBs (MOI 10) were added to cells that had been incubated with DMSO (0,1%) or Taxol (10 µM) via centrifugation at 4°C followed by fixation. n=3 representing 30 cells each, Error bars denote ±SEM, ns; not significant (two-tailed unpaired Student´s *t*-test).

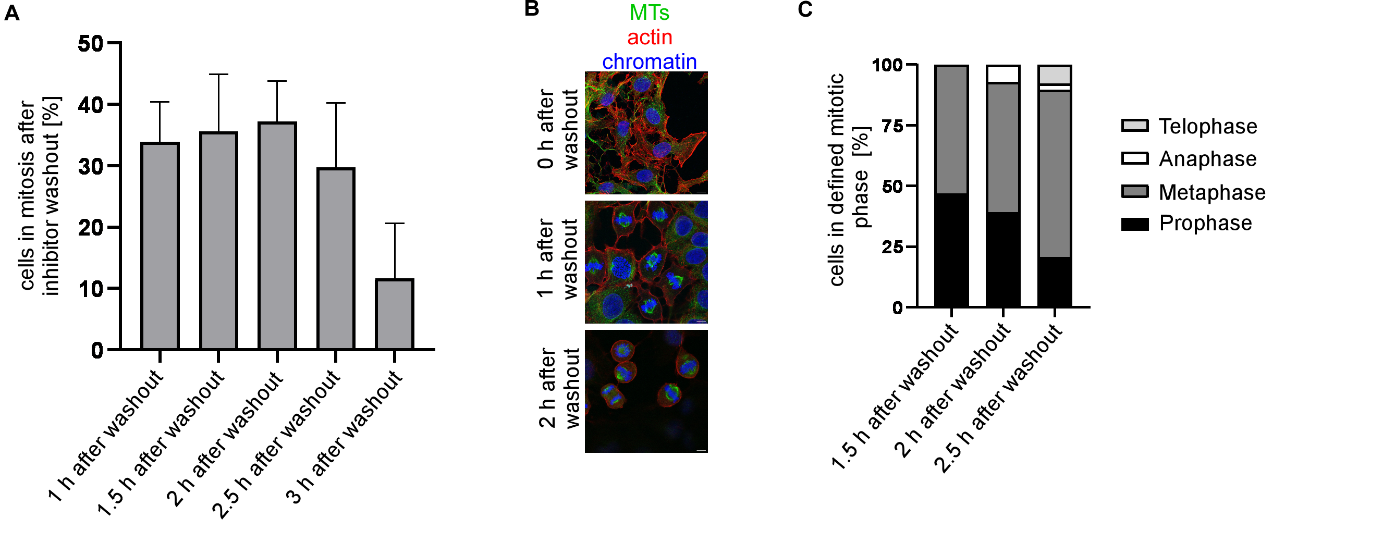

**Fig S3. RO-3306 inhibitor treatment of mammalian cells**

**A** Quantification of the number of mitotic cells after 1 to 3 h of RO-3306 washout. Cells were arrested at G2/M by treatment with RO-3306 for 20 h. Cells were defined as mitotic cell when in any mitotic spindle stage. n=3 each representing 80 cells. Error bars denote ±SEM. **B** Representative confocal fluorescence images of mitotic U2OS cells treated with and released from CDK1 Inhibitor RO-3306 at indicated time points. MTs were visualized with anti-α-tubulin antibody (green), actin with rhodamine-phalloidin staining (red) and DNA with DAPI (blue). Scale bars: 10 µm. **C** Microscopic determination and quantification of various mitotic stages 1.5 to 2.5 h after RO-3306 washout. Mitotic stage was determined based on spindle structure. n=3 at least 20 mitotic cells for each time point.

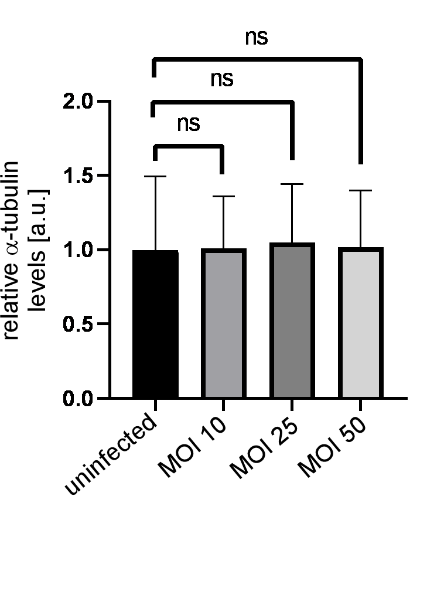

**Fig S4. Relative α-tubulin levels in cells infected with different MOIs**

Quantification of relative α-tubulin levels 1 hpi. Band intensities from Western blots of α-tubulin and GAPDH in cells infected with different MOIs of *C. pneumoniae* EBs were determined using ImageJ. The ratio of α-tubulin levels to GAPDH levels was analysed and compared relative to control cells (The ratio for uninfected control set to 1). n=3 independent experiments/condition, Error bars denote ±SEM, ns; not significant (two-tailed unpaired Student´s *t*-test).

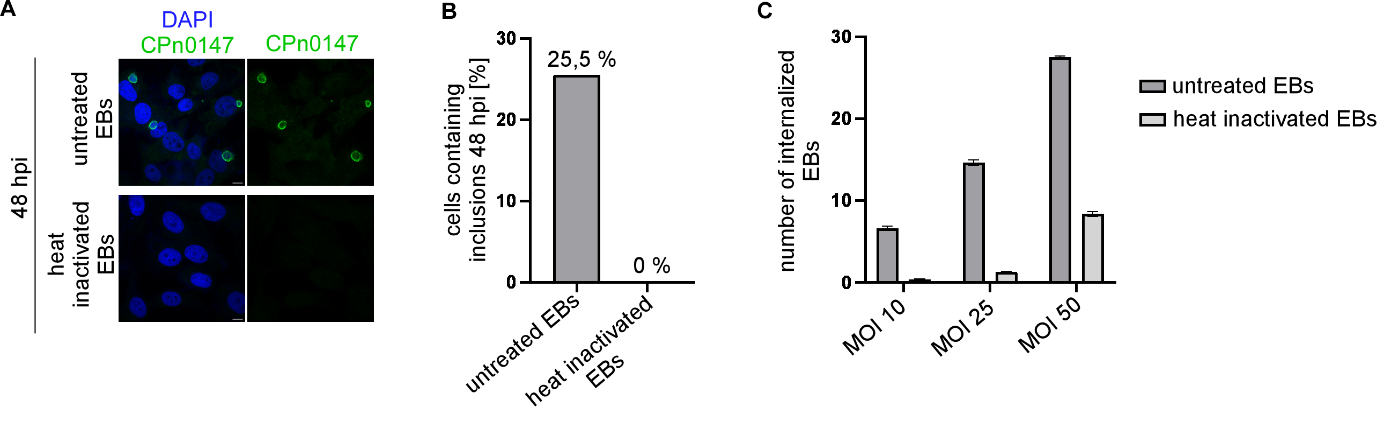

**Fig S5. Heat-inactivated EBs and infection capability**

**A** Representative confocal image of U2OS cells incubated with untreated or heat-inactivated *C. pneumoniae* EBs (MOI 5) for 48 h. Inclusion membrane was visualized via anti-CPn0147 antibody (chlamydial Inc protein) in green and DNA with DAPI (blue). Images shown are maximum intensity projections. Scale bars: 10 µM. **B** Quantification of the number of infected cells using untreated or heat-inactivated EBs. Data obtained 48 hpi. n=100 cells per condition. **C** Quantification of the number of internalized EBs in cells infected with untreated or heat inactivated *C. pneumoniae* EBs at the indicated MOI for 1 h. n=3 each representing 20 cells per condition. Error bars denote ±SEM.

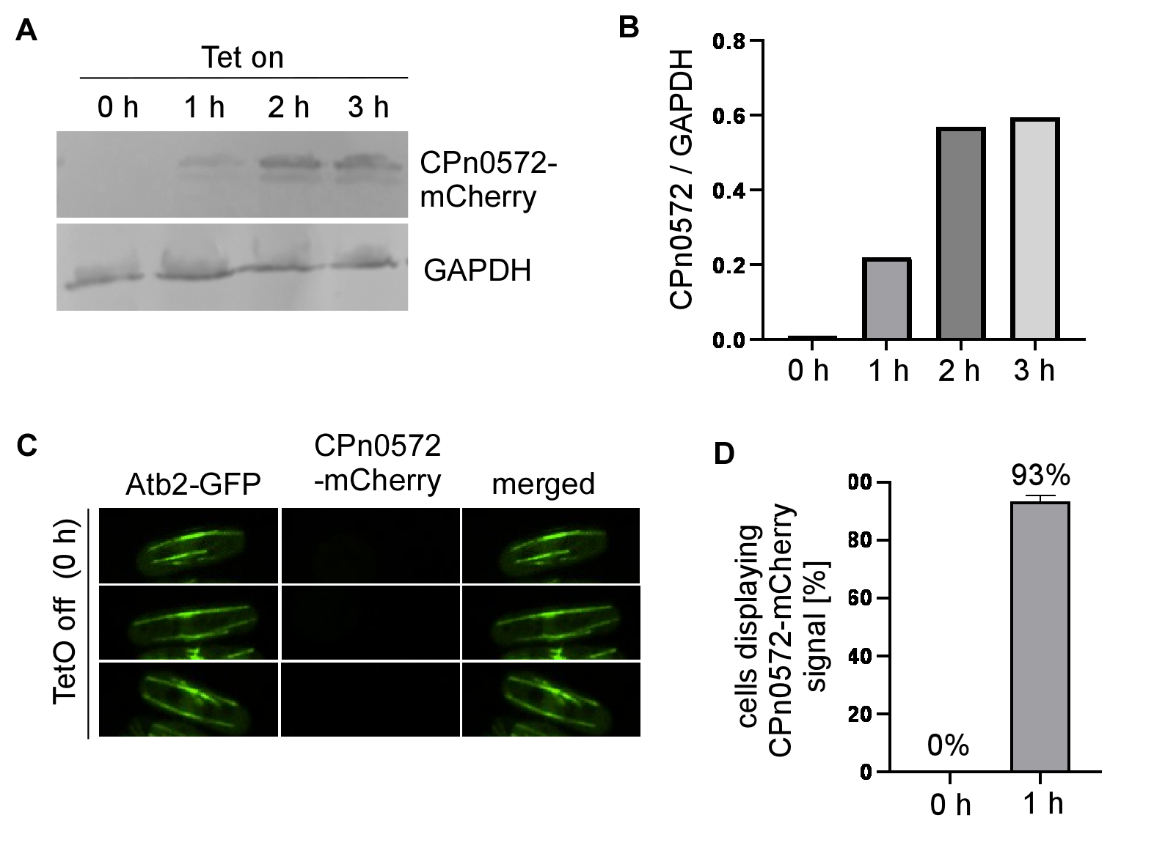

**Fig S6. Expression of genome integrated CPn0572 under control of TetO promotor in *S. pombe***

**A** Western blot analysis of CPn0572-mCherry expression under control of the TetO promotor in *S. pombe*. Logarithmically growing cells were incubated at 30°C with 2,5 µg tetracycline for the indicated time points followed by cell lysis. Protein extracts were separated on a 10% SDS-PAGE. Western blot analysis was carried out using anti-mCherry antibody and GAPDH antibody. **B** Quantification of the amount of CPn0572-mCherry versus GAPDH shown in **A** using ImageJ. **C** Confocal images of the α-tubulin-GFP *S. pombe* strain containing genome-integrated *CPn0572-mCherry* grown at 30 °C under TetO promoter-off conditions (no tetracycline). MTs were visualized in living cells via α-tubulin-GFP (Atb2-GFP). Images shown are maximum intensity projection. **D** Quantification of the number of cells displaying CPn0572-mCherry signals at 0 and 1h after induction with 2,5 µg tetracycline at 30°C. n=7 each containing at least 22 cells per time point.

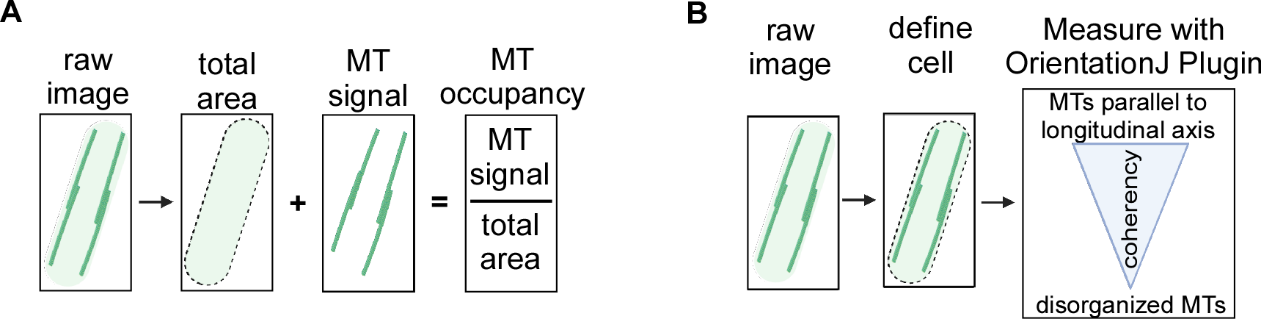

**Fig S7. Quantification methods of spatial MT distribution using ImageJ**

**A** Schematic representation of the analysis to determine MT occupancy i.e. the area in the cell occupied by α-tubulin signal. ImageJ was used to measure the MT signal in relation to the total cell area (MT occupancy). **B** Schematic representation of quantification of MT orientation along the long axis using the OrientationJ Plugin of ImageJ. A high MT coherency indicates that MTs are oriented parallel to the longitudinal axis of the cell while a low MT coherency represents disorganized MTs.

**Supplementary movie legends**

**Movie 1** Live cell images of wild type α-tubulin-GFP expressing *S. pombe* cells. 11,3 seconds interval, total 56,5 seconds. Scale bar: 5 µm.

**Movie 2** Live cell images of α-tubulin-GFP and CPn0572-mCherry expressing *S. pombe* cells. 11,3 seconds interval, total 79,1 seconds. Scale bar: 5 µm.
